# Hippocampal and prefrontal contributions to memory-guided navigation depend on task epoch

**DOI:** 10.64898/2026.05.29.728861

**Authors:** Tora Dohi, Lucy Anderson, György Buzsàki, Ipshita Zutshi

**Affiliations:** Neuroscience Institute, New York University Grossman School of Medicine; Department of Neuroscience and Cell Biology, Rutgers Robert Wood Johnson Medical School

**Keywords:** Hippocampus, working memory, perturbation, prefrontal cortex, optogenetics

## Abstract

The hippocampus and medial prefrontal cortex (mPFC) are required for delayed working-memory tasks, but when in the task their engagement becomes necessary remains an open question. We trained mice on a delayed cue-guided T-maze navigation task and used transient optogenetic silencing to test the contribution of the mPFC and hippocampus during distinct task epochs. Surprisingly, silencing during the delay period did not impair performance. In contrast, perturbation during the early phase of the central arm traversal produced robust deficits. Mice made more perseverative choices, slowed their speed and ran farther down the track before turning towards the selected arm. Control experiments showed that these effects could not be explained by the manipulation targeting elapsed time and were rather specific to task phase. These results indicate that hippocampal and prefrontal contributions to memory-guided behavior are not uniform across the trial, but instead depend on task epoch. More broadly, they suggest that the functional engagement of these circuits is gated by behavioral state or context.

## INTRODUCTION

Working memory tasks with delayed responses consistently implicate an essential role of the hippocampus and medial prefrontal cortex (mPFC) areas. Lesions, pharmacological inactivation, and chronic perturbations of either region disrupt performance across species (Aggleton et al., 1986; Ainge et al., 2007; Dudchenko et al., 2000; Eichenbaum, 2017; Olton et al., 1979; Sabariego et al., 2019; Scoville and Milner, 1957; Squire and Zola-Morgan, 1991).

Despite this clear necessity, hippocampal and prefrontal contributions to memory-guided behavior are typically inferred over relatively coarse timescales, leaving unresolved when within a trial these regions are causally required. Because memory-guided decisions unfold on short timescales, resolving the timing of circuit engagement may determine how stored information is translated into behavior.

A prevailing view is that activity in the hippocampus bridges temporally disconnected events, linking past cues to future choices, while the mPFC maintains task-relevant reference frames or schemas across delays (Eichenbaum, 2017, 2014; MacDonald et al., 2011; Muysers et al., 2024; Pastalkova et al., 2008; Taxidis et al., 2020). Yet activity in these regions during delayed tasks is often sparse, variable, and distributed across networks , and in many cases is only weakly related to behavioral performance (Choong Yong et al., 2022; Sabariego et al., 2019; Yuan et al., 2025). These observations raise the possibility that hippocampal and prefrontal contributions are not persistent across a trial, but instead are engaged at specific moments during memory-guided behavior.

To determine when hippocampal and prefrontal activity is causally required during a memory-guided navigation task, we developed a delayed cue-guided paradigm in which each brain region could be transiently silenced at distinct task epochs. This approach allowed us to test whether the contribution of these regions can be uncovered based on the timing of disruption. Using this design, we find that hippocampal and mPFC contributions to behavior are temporally specific, with perturbations during central arm traversal producing robust impairments, whereas manipulations during other epochs having minimal effects.

## RESULTS

### A delayed cue-guided navigation task with temporally structured trial epochs

To assess how the hippocampus and mPFC contribute to memory-guided behaviors, we trained mice (n = 8) on a delayed cue-guided navigation task, conceptually similar to previously described cue-based navigation tasks (Ainge et al., 2012; Harvey et al., 2012; Muysers et al., 2024; Sun et al., 2023; Taxidis et al., 2020), but with more temporally structured trial epochs (**Fig. 1a**). Each trial began with the mouse initiating a nose-poke at a door at the home base, which triggered a 1-second-long lateralized visual cue (LED strip on either the left or right wall of the central stem), followed by a 1-second delay period behind the door. At the end of the delay, the door opened, and the animal ran down a 1-meter-long track and turned into the arm corresponding to the previously cued side. The median duration of this *Run* phase was 2.41s (**Fig. 1b**). Correct choices were rewarded at the end of the selected arm, after which the mouse returned to the home base to receive a second water reward and subsequently initiate the next trial. Trained mice achieved ∼80% accuracy in the task. Examination of left- and right-ward trajectories showed divergence only in the final segment of the track (**Fig. 1c, d**), indicating that animals did not adopt an early motor strategy (e.g., immediately biasing one side) and instead relied on the remembered cue to guide their choice. Importantly, the location of trajectory divergence presumably reflects when the decision is expressed in behavior, not necessarily when it is formed.

**Figure 1.**
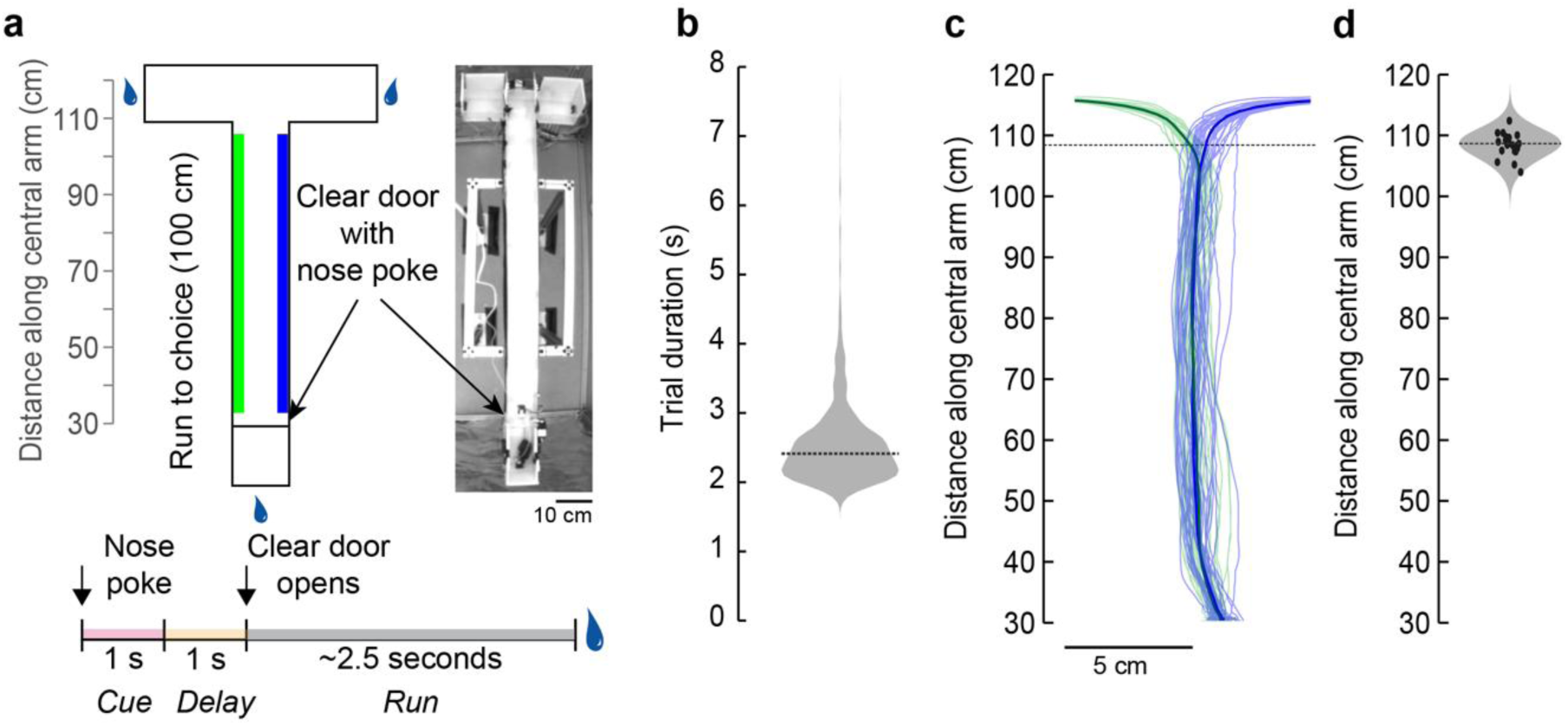
Task design and behavioral trajectories in a delayed cue-guided navigation task. (a) *Left*, mice initiated each trial by nose-poking at a clear door at the home base. The nose-poke triggered a 1-second visual cue where an LED strip on either the right (*blue*) or left (*green*) wall turned on. This was followed by a 1-second delay period with no cues. At the end of the delay period, the door opened, and the mice must run the length of the track (1 m) and turn toward the side of the arm where the cue was presented. Trials were reset when the mice returned to the water port at the home base. Incorrect trials were not rewarded. *Right,* top view of the track. (b) Average duration of the *Run* epoch of a trial (n = 1517 trials from 8 mice). (c) Example session showing left-ward (*green*) and right-ward (*blue*) trajectories. Thick lines show the mean trajectory. Only correct trials were included. *Dashed horizontal line,* position in the central arm where left-ward and right-ward trajectories diverged significantly. (d) Distribution of the position of significant divergence between left-bound and right-bound trials across all sessions (n = 27 sessions from 2 mice). The absence of early divergence indicates that trajectories did not differ at trial onset.

To test the role of the hippocampus and mPFC in different phases of the task, we used optogenetic inhibition of pyramidal neurons by activating either Parvalbumin-positive (PV+) interneurons or Dlx-expressing (Dimidschstein et al., 2016) inhibitory neurons to achieve widespread suppression of spiking in pyramidal cells (Fernández-Ruiz et al., 2021; Olsen et al., 2012; Zutshi et al., 2022). Five mice expressed ChR2 in PV+ interneurons (PV-Cre × Ai32), while two mice were injected with AAV-Dlx-ChR2 in both the dorsal hippocampus (CA1) and prelimbic mPFC (**Fig. 2**). One additional wild-type mouse served as a sham-stimulation control. Optic fibers were bilaterally implanted in the hippocampus and/or mPFC, depending on the experimental group, and histology confirmed targeting and viral expression in all animals (**Fig. 2**).

**Figure 2.**
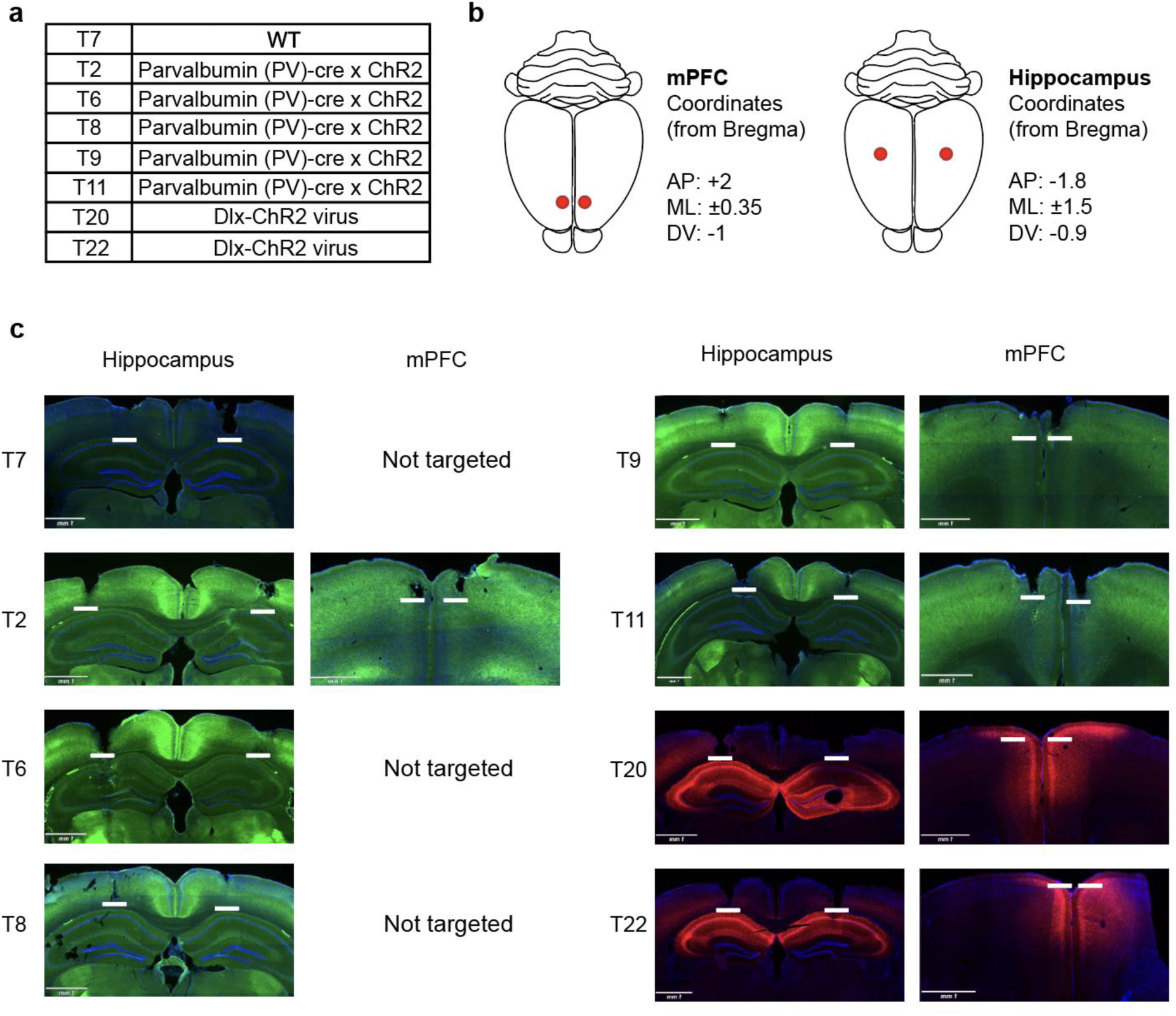
Strategy to optogenetically silence dorsal hippocampus (CA1) and prelimbic mPFC. (a) Experimental animals (n = 8, including one wild-type control). Five mice were Parvalbumin (PV)-cre x Ai32 mice, two mice were injected with an AAV expressing Dlx-ChR2 in the dorsal hippocampus (CA1) and prelimbic mPFC. (b) Stereotaxic coordinates targeting mPFC and dorsal hippocampus. (c) Histological verification of viral expression and fiber placement for each mouse. T7, T6, and T8 had bilateral hippocampal fibers only; the remaining mice had bilateral fibers in both the hippocampus and mPFC. *Green*, ChR2-GFP expression in PV+ cells. *Red*, ChR2-mCherry expression in Dlx+ cells. *Horizontal white lines* correspond to optic fiber tips. *Scale bar,* 1 mm.

**Figure 3.**
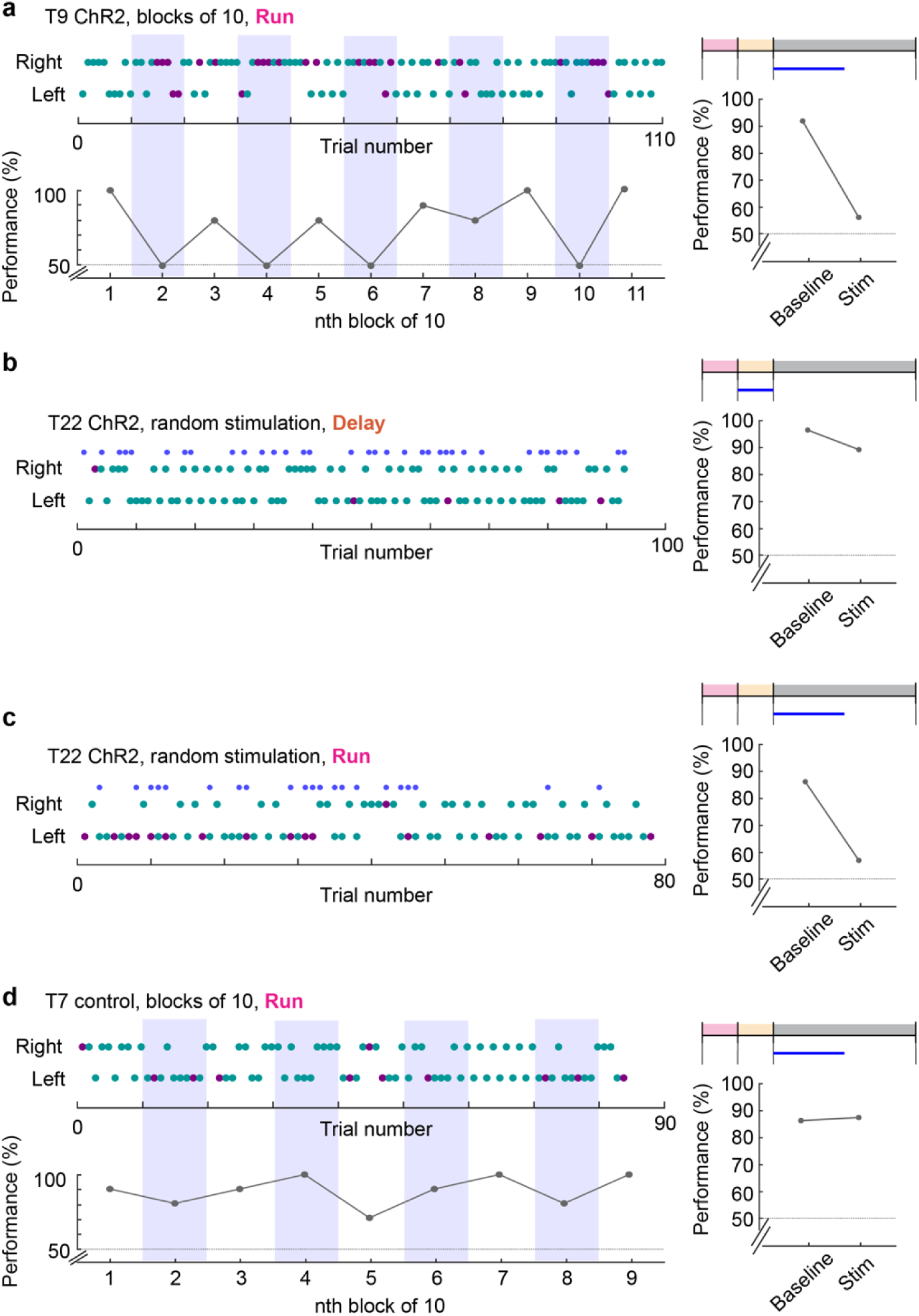
Example sessions illustrating epoch-specific effects of optogenetic silencing. (a) Example session from a PV-Cre x ChR2 mouse (T9) with hippocampal silencing during the early run phase (first 2s). *Top-left*, trial-by-trial choices (*green*, correct; m*agenta*, incorrect, with shaded blocks indicating stimulation trials). *Bottom-left,* average performance in blocks of 10 trials. *Top-right,* schematic of the stimulated task-epoch. *Bottom-right,* average performance during baseline and stimulation trials. (b) Example session from a Dlx-ChR2 mouse (T22) with silencing during the delay period (∼30% of trials, randomly interleaved). Format as in (a). Block-averaged performance is not shown as each block contains a mixture of stimulation and baseline trials. (c) Same mouse as in (b) (T22), with silencing during the early run phase (∼30% of trials). (d) Example session from a control mouse (T7) with light delivery during the early run phase. No impairment is observed.

During behavioral sessions, optogenetic stimulation was delivered in either blocks of 10 consecutive trials or on a random 30% subset of trials. Light was applied during one of nine trial epochs (Cue, Delay, Cue+Delay, Run (early), Run (mid), Run (late), Delay+Run, Longer Delay, and Unilateral stimulation), allowing us to test how the effects of perturbation depend on the timing of disruption within the trial (**Fig. 4a, 5a**).

**Figure 4.**
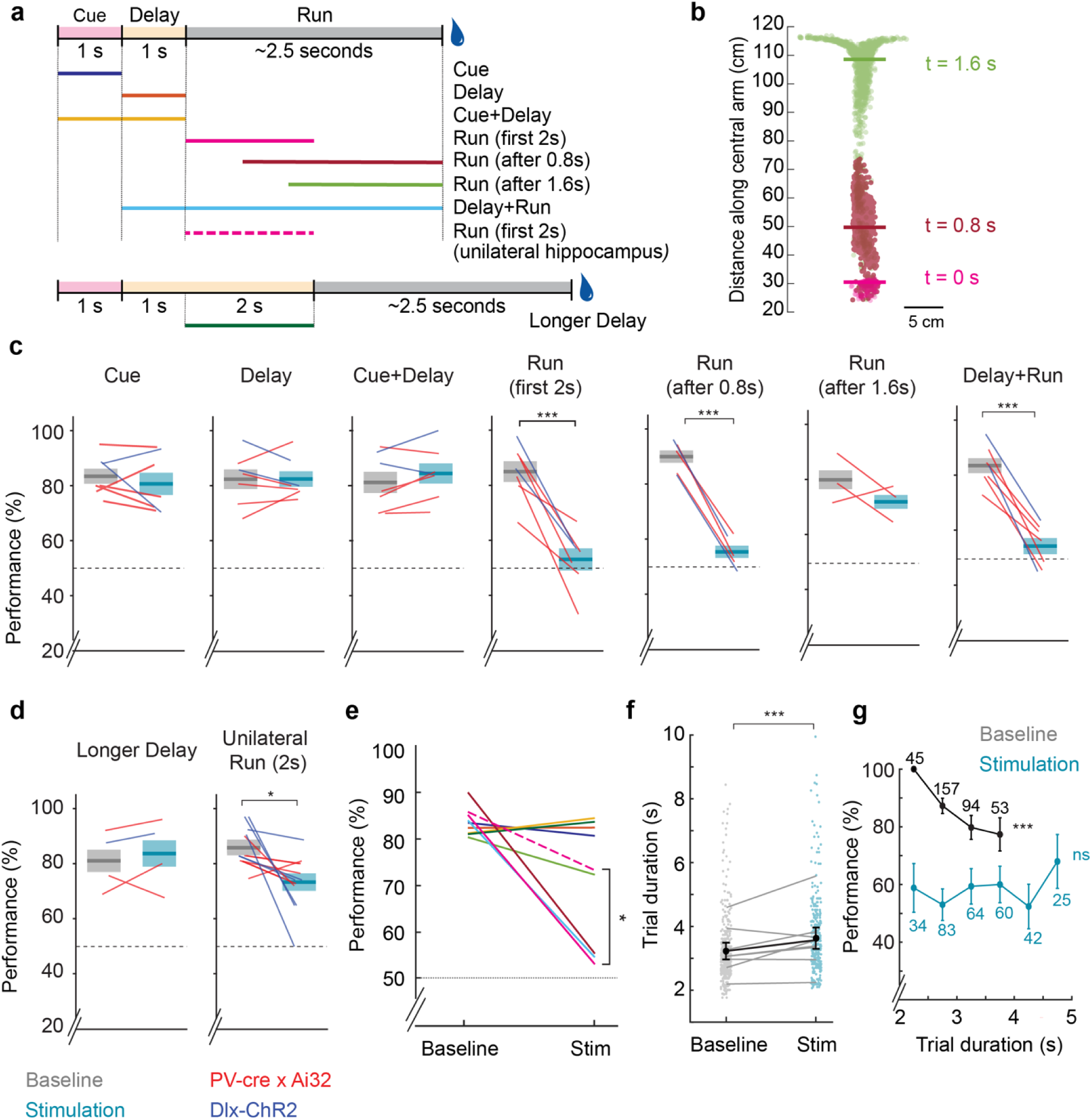
Optogenetic silencing of dorsal hippocampus reveals epoch-specific effects on memory-guided behavior. (a) Task epochs targeted for optogenetic stimulation. To control for time since cue delivery, a subset of mice was tested with a longer delay, with stimulation applied to the final 2 s of the delay. Unless otherwise noted, all manipulations were bilateral. (b) Position of the animal along the central arm at the onset of stimulation for the three run manipulations. Stimulation windows were aligned to distinct portions of the run epoch based on median trial duration (∼2.4 s): immediately after door opening (t = 0 s), after the stop-to-run transition had occurred (t = 0.8 s), and near the choice point (t = 1.6 s). Each dot represents the position from a single trial. Horizontal lines indicate the median position at those timepoints. (c, d) Performance during baseline (gray) and hippocampal stimulation (blue) trials across task epochs. Each line represents the average performance of one mouse. Red lines, PV-Cre × Ai32 mice; blue lines, Dlx-ChR2 mice. Perturbations during early central arm traversal produced large and consistent impairments across animals, whereas manipulations during other task epochs resulted in minimal effects. (Paired t-test; Cue: n = 7 mice, t(6) = 0.8177, *p* = 0.4448; Delay: n = 7 mice, t(6) = -0.0218, *p* = 0.9833; Cue+Delay: n = 7 mice, t(6) = -1.6816, *p =* 0.1436; Run (first 2s): n = 7 mice, t(6) = 6.6353, *p =* 5.653 × 10^−4^; Run (after 800 ms): n = 5 mice, t(4) = 19.797, *p* = 3.841 × 10^−5^; Run (after 1.6s) : n = 3 mice, t(2) = 1.1407, *p =* 0.3722; Delay + Run: n = 7 mice, t(6) = 8.2827, *p* = 1.677 × 10^−4^; Longer delay: n = 4 mice, t(3) = -0.6665, *p* = 0.5527; Unilateral hippocampus run (first 2s): n = 9 mice, t(8) = 2.4801, *p =* 0.0381). (e) Summary of the average effect across stimulation conditions. Colors correspond to stimulation windows shown in (a). The largest impairments were observed for manipulations targeting early central arm traversal, whereas other conditions showed smaller or inconsistent effects (Comparison between Run (first 2s), bilateral and unilateral hippocampus, t-test of difference from baseline, n = 9 sessions from 7 mice, t(14) = 2.7326, *p = 0.0162*). (f) Trial duration during baseline and stimulation trials for the bilateral hippocampal Run (first 2 s) manipulation. Hippocampal silencing increased trial duration. Each dot is a trial. *Gray lines* connect the mean trial duration for individual mice across conditions (Wilcoxon rank sum test, n = 414 & 338 trials, Z = -5.64, *p = 1.68 × 10^−8^*). (g) Relationship between trial duration and performance during baseline and hippocampal stimulation trials. During baseline trials (*black*), slower trials were associated with reduced performance (Spearman correlation, R = -0.21, *p = 1.37 × 10^−5^*). In contrast, during hippocampal silencing (*blue*), this effect of trial duration was lost with uniform impairment (Spearman correlation, R = - 0.01, *p = 0.787*). Numbers indicate the number of trials contributing to each bin. *\*p<0.05, ***p<0.001*

**Figure 5.**
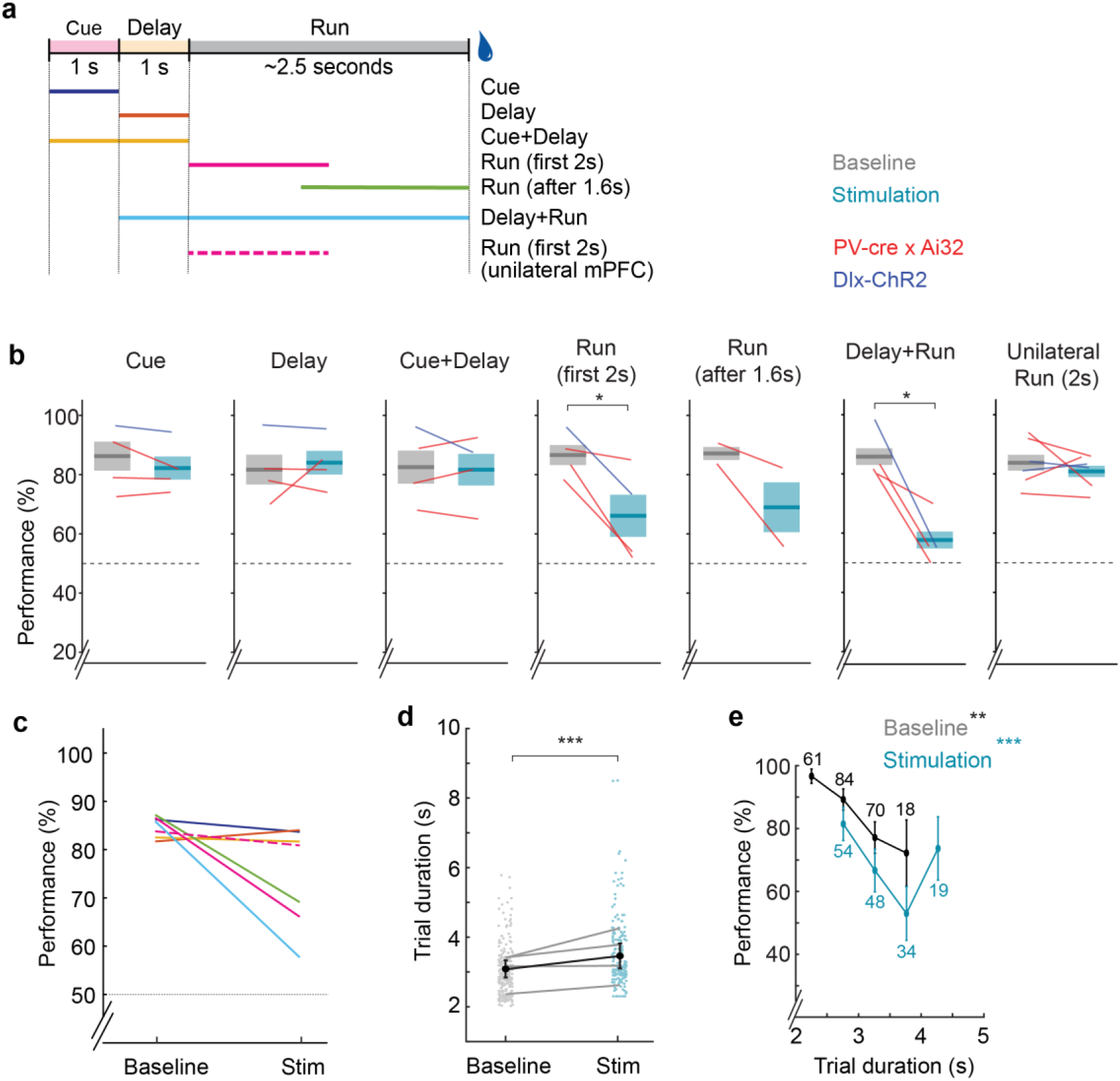
Optogenetic silencing of prelimbic mPFC leads to epoch-specific effects on memory-guided behavior. (a) As in Fig. 4a, schematic showing the task epochs targeted for mPFC silencing. (b) Average performance of baseline (*gray*) versus mPFC stimulation (*blue*) trials across mice. Each line corresponds to the average performance of a single mouse. *Red lines*, PV-Cre x Ai32 mice. *Blue lines*, Dlx-ChR2 virus-injected mice. Perturbations targeting early central arm traversal produced impairments, whereas manipulations during other task epochs resulted in smaller or inconsistent effects. (Paired t-Test; Cue: n = 4 mice, t(3) = 1.098, *p =* 0.3524; Delay: n = 4 mice, t(3) = -0.5466, *p* = 0.6227; Cue+Delay: n = 4 mice, t(3) = 0.2722, *p =* 0.8031; Run (first 2s): n = 4 mice, t(3) = 3.4740, *p* = 0.0402; Run (after 1.6s): n = 2 mice, t(1) = 1.8526, *p =* 0.3151; Delay + Run: n = 4 mice, t(3) = 4.09, *p* = 0.0264; Unilateral mPFC run (first 2s): n = 6 mice, t(5) = 0.8203, *p =* 0.4494). (c) Summary of the average effect from (b), comparing across stimulation conditions. Colors correspond to stimulation windows shown in (a). As with hippocampal silencing, only stimulations targeted to the central arm traversal led to an impairment. Unilateral manipulations had very little effect. (d) Trial duration during baseline and stimulation trials for the bilateral mPFC Run (first 2 s) manipulation. Gray lines connect the mean trial duration for individual mice across conditions (Wilcoxon rank sum test, n = 257 & 188 trials, Z = -6.2, *p = 5.42 × 10^−10^*). (e) Relationship between trial duration and performance during baseline and mPFC stimulation trials. Same as Fig. 4g. Baseline trials (*black*) (Spearman correlation, R = -0.17, *p = 0.0067*). The effect of trial duration persisted during mPFC silencing (Spearman correlation, R = -0.32, *p = 1.02 × 10^−5^*). Numbers indicate the number of trials contributing to each bin. \**p*<0.05, \*\**p*<0.01, \*\**\*p*<0.001

### Hippocampal and mPFC silencing produces epoch-specific impairments in memory-guided behavior

Mice showed consistent and reversible impairments when hippocampal silencing was targeted to the early portion of the Run epoch (**Fig. 3a**). Performance decreased during stimulation trials and returned to baseline immediately afterward. Similar effects were observed in Dlx-ChR2 virus-injected mice when silencing occurred during this same epoch (**Fig. 3c**). This effect was consistent across sessions where stimulation was delivered in blocks versus random stimulation.

Perturbations during other task epochs, including the cue and delay periods produced smaller or inconsistent effects (**Fig. 3b, 4c, 4e**). Light delivery in a control mouse during the early Run epoch did not affect performance (**Fig. 3d**), indicating that the observed deficits were not due to light stimulation alone.

To examine whether the effect of perturbation varied across different stages of central arm traversal, we next targeted sub-portions of the Run epoch (**Fig. 4a**). Based on a median Run length of 2.4 s, stimulation was delivered either at run initiation (first 2s, starting at 0 s), after the mice had already started running (starting at 0.8 s), or near the choice point (starting at 1.6 s) (**Fig. 4b**). Bilateral hippocampal silencing produced the largest and most consistent impairments when stimulation occurred during the early and middle portions of the run (**Fig. 4c**). In contrast, perturbations near the choice point produced smaller and more variable effects across animals. Thus, the effects of perturbation were specific to the early portion of central arm traversal, after movement initiation but before reaching the choice point, suggesting that hippocampal activity is particularly important during active navigation before behavioral commitment to the selected arm.

Bilateral hippocampal silencing during the Run epoch increased trial duration, indicating that mice slowed during stimulation (**Fig. 4f**). Under baseline conditions, slower trials were associated with poorer performance (**Fig. 4g**). During hippocampal silencing, however, performance was impaired across trial durations (**Fig. 4g**), suggesting that the deficits could not be explained solely by slower movement or prolonged trial times.

To test whether the effect of hippocampal perturbation depended on time elapsed from cue presentation rather than task epoch, we trained a subset of mice on a longer delay (3 s) condition. In this version of the task, stimulation was applied to the final 2 s of the delay, matching the early Run epoch in terms of elapsed time from cue presentation. Under these conditions, silencing during the delay did not reproduce the strong impairments observed during early central arm traversal (**Fig. 4d**). Although this cohort was smaller, the effect sizes were minimal and inconsistent across animals, in contrast to the large and reliable deficits observed during run-phase perturbations. These results argue against a simple time-from-cue explanation and instead indicate that the impact of hippocampal perturbation depends on task epoch.

Finally, unilateral hippocampal silencing during early central arm traversal produced a modest impairment that was smaller than bilateral effects (**Fig**. **4d, e**), suggesting partial compensation across hemispheres.

We next tested whether mPFC perturbation produced a similar pattern of effects across task epochs. As with hippocampal manipulations, stimulation was targeted to distinct epochs within a trial (**Fig. 5a**). Similar to hippocampal perturbation, bilateral mPFC silencing during the early central arm traversal and during the combined Delay+Run condition resulted in significant impairments (**Fig. 5b**). In contrast to hippocampal perturbations, unilateral mPFC silencing did not produce a measurable impairment (**Fig. 5b–c**). Bilateral mPFC silencing during the Run epoch also increased trial duration, indicating that mice slowed during stimulation (**Fig. 5d**).

Unlike the effect of hippocampal silencing however, slower trials were associated with poorer performance in both baseline and stimulation trials (**Fig. 5e**). Thus, the relationship between trial duration and performance was preserved during mPFC perturbation, despite an overall decrease in performance. These results suggest that the behavioral impairments produced by hippocampal and mPFC silencing may arise from distinct underlying mechanisms.

### Stimulation induces behavioral perseveration and alters trajectory dynamics

To further characterize how early run-phase silencing disrupts behavior, we analyzed trial-by-trial arm choices and movement trajectories. Both hippocampal and mPFC silencing during early central arm traversal increased the likelihood of repeated same-arm choices across consecutive trials (**Fig. 6a–c**), consistent with perseverative behavior. This shift was not explained by biases in cue delivery (**Fig. 6b**).

**Figure 6.**
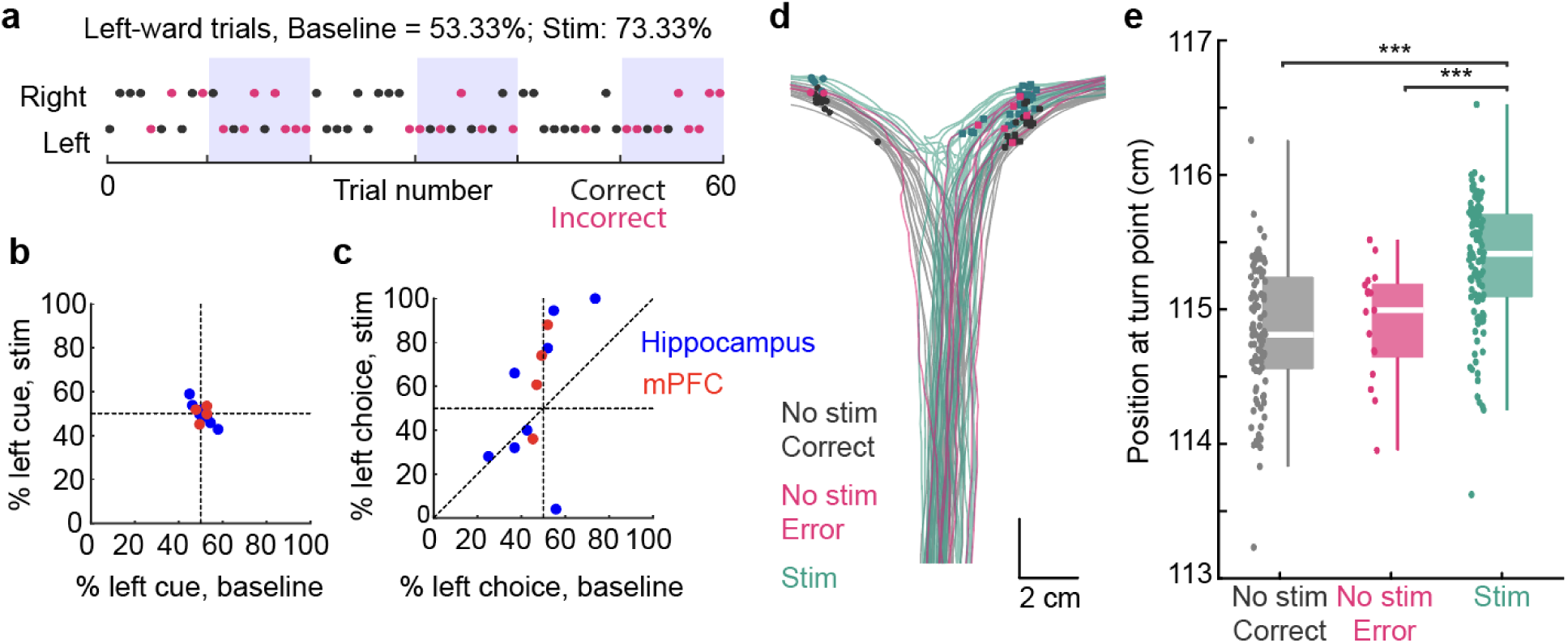
Optogenetic silencing during early central arm traversal increases perseveration and alters behavioral trajectories. (a) Example session where optogenetic stimulation targeted hippocampal fibers during *RUN* (first 2s). Trial-by-trial behavior is shown with leftward and rightward choices. *Black*, correct trials; *magenta*, incorrect trials; blue boxes, blocks of 10 stimulation trials. The mouse chose the left arm on 73.33% of stimulation trials compared to 53.33% during baseline, consistent with a leftward bias during stimulation. (b) Percentage of leftward cues during baseline vs. stimulation trials. Across sessions, cue distribution remained near 50% for both hippocampus (*blue*) and mPFC (*red*) targeting. (c) Same as (b), but for the **chosen** arm rather than the cue. While baseline choices were near 50%, stimulation induced a directional bias (*left bias*, above horizontal line; *right bias*, below horizontal line). (d) Example trajectories from a hippocampal stimulation session targeting the *RUN* phase. *Gray*, correct no-stimulation trials; *magenta*, incorrect no-stimulation trials; *teal*, stimulation trials (correct and incorrect). Circles mark leftward turns; squares mark rightward turns, defined as the point of maximal horizontal motion into the side arms. (e) Position along the central arm at the point of maximal horizontal motion toward a side arm. *Gray*, correct no-stimulation trials; *magenta,* incorrect no-stimulation trials; *teal*, stimulation trials (correct and incorrect). Each dot is a trial. During stimulation, but not during no-stimulation error trials, the point of maximal horizontal motion occurred farther along the central arm (Kruskal Wallis test followed by Tukey-Kramer post-hoc tests, n = 98,17,110 trials respectively, Chi-sq (2) = 56.601, *p = 5.12 × 10^−13^).* All box plots show median ± interquartile; whiskers show range excluding outliers.*\*\*\*p<0.001*.

Examination of movement trajectories revealed that, during stimulation, mice advanced farther along the central arm before initiating a turn (**Fig. 6d, e**). This delayed commitment was not observed during no-stimulation error trials. These findings imply that hippocampal or mPFC perturbation during early central arm traversal biases action selection and alters the dynamic of movement to potential habitual or motor-biased strategies (Packard and McGaugh, 1996).

## DISCUSSION

By transiently silencing hippocampal and mPFC neurons at defined task epochs, we found that the impact of perturbation depends on when within the trial activity is disrupted. Surprisingly, silencing during the cue or delay periods produced little to no impairment, despite these epochs being the canonical locus of working memory encoding and maintenance. Behavioral impairments were instead largest and most consistent when either region was silenced during early central arm traversal. Matching stimulation during a longer delay did not reproduce these deficits, arguing against a simple dependence on time since cue presentation. Together, these results indicate that hippocampal and prefrontal contributions to memory-guided behavior are not uniform across the trial, but depend on task epoch, and more specifically, on whether the animal is actively navigating toward a decision.

### Single-region versus distributed memory models

Traditionally, the hippocampus and mPFC have been viewed as core substrates for working memory. Lesion and inactivation studies show that damage to either structure disrupts delayed memory tasks, including delayed spatial alternation, while continuous versions of the same tasks remain intact (Aggleton et al., 1986; Ainge et al., 2007; Dudchenko et al., 2000; Olton et al., 1979; Sabariego et al., 2019). Patients with hippocampal–prefrontal lesions similarly exhibit profound deficits in working memory (Scoville and Milner, 1957; Squire and Zola-Morgan, 1991). At the neural level, reports of sequential activity during the delay period in the hippocampus (Baeg et al., 2003; Eichenbaum, 2014; Fujisawa et al., 2008; MacDonald et al., 2011; Pastalkova et al., 2008) or persistent activity at the single neuron and population level in the PFC (Constantinidis et al., 2002; Funahashi, 2006; Fuster and Alexander, 1971; Kubota and Niki, 1971; Miller et al., 1996; Murray et al., 2017; Procyk and Goldman-Rakic, 2006), reinforced the idea that working memory is actively maintained by delay-period firing within these regions.

However, more recent large-scale recordings have begun to modify this notion. Delay-period activity appears distributed across multiple regions, with loop-based dynamics across interconnected areas sustaining memory (Lundqvist et al., 2018; Voitov and Mrsic-Flogel, 2022). Hippocampal delay sequences themselves are brief, variable, and not predictive of behavioral performance (Choong Yong et al., 2022; Sabariego et al., 2019; Yuan et al., 2025). These findings collectively support a distributed-memory framework in which representations are maintained across interacting cortical and subcortical nodes rather than localized to any one anatomical structure (Christophel et al., 2017; Courtney, 2022; Guo et al., 2014; Slotnick, 2023, 2022; Stern and Hasselmo, 2022; Stokes, 2015). . Under this framework, memory may persist through activity-silent mechanisms, such as short-term synaptic facilitation, that are resilient to transient suppression of spiking within a single region (Stokes, 2015).

Our results are consistent with this framework. Silencing the hippocampus or mPFC during the delay produced little impairment, suggesting that whatever representation bridges cue and choice does not depend on sustained spiking within either region in isolation. This is consistent with a scenario in which memory is maintained through activity-silent or distributed mechanisms during the delay, and is only re-engaged, and therefore vulnerable to perturbation, when the animal initiates active navigation toward a decision.

### Hippocampal and mPFC contributions are temporally specific within the trial

Longer delays are often expected to place greater demands on hippocampal-dependent memory (Squire and Zola-Morgan, 1991), and our own behavioral data are consistent with this idea: under baseline conditions, performance declined when mice took longer to run down the central arm. However, several observations argue that the effects of hippocampal silencing cannot be explained simply by the passage of time from cue presentation. Silencing during a longer delay did not reproduce the strong impairment observed during early central arm traversal, even when stimulation was matched for elapsed time from the cue. In addition, hippocampal silencing during the Run epoch degraded performance across trial durations, abolishing the normal relationship between trial duration and accuracy. This contrasts with mPFC silencing, where performance was impaired but the relationship between trial duration and accuracy was preserved. Together, these dissociations suggest that hippocampal and mPFC contributions during active navigation are distinct, and that the hippocampal effect cannot be reduced to a general slowing or elapsed-time account.

Behaviorally, these impairments were accompanied by slower movement, delayed choice commitment, and increased perseverative responding. Animals frequently continued farther down the track before committing to a turn, and showed a directional bias toward the previously chosen arm — a pattern consistent with a shift toward response-based or habitual strategies (Packard and Goodman, 2012; Packard and McGaugh, 1996). Thus, hippocampal and mPFC perturbation during early navigation did not simply impair memory retrieval, but altered the balance between flexible memory-guided behavior and more automatic response selection.

Our results can be compared with studies that have manipulated hippocampal–mPFC circuits during distinct phases of delayed non-match-to-sample tasks. Spellman et al. selectively silenced ventral hippocampal inputs to mPFC and observed impairments when perturbations were delivered during encoding, but not during delay or retrieval (Spellman et al., 2015).

Importantly, the encoding phase in their task involved active navigation down the maze arm, not passive cue observation. Babl and Sigurdsson (2025) similarly reported that both dorsal and ventral hippocampus contributed during the sample phase of a spatial working memory task, whereas only dorsal hippocampus was necessary during the choice phase (Babl and Sigurdsson, 2025). Studies manipulating nucleus reuniens circuits have reported impairments during either encoding or retrieval depending on task structure (Maisson et al., 2018; Rahman et al., 2021), further suggesting that no single phase is universally privileged. Across these studies, a common pattern emerges: perturbations are most disruptive when delivered during epochs that require active behavioral engagement, and relatively ineffective during passive maintenance periods. This suggests that hippocampal–mPFC contributions may not map cleanly onto predefined phase labels such as encoding, delay, or retrieval, but instead track whether the animal is in an active, navigation-engaged state.

A key difference in how hippocampal circuits are engaged across task epochs may relate to the specific theta states present during behavior. Hippocampal theta oscillations are commonly divided into two broad classes: Type I theta, which is atropine-resistant and prominent during locomotion and active exploration, and Type II theta, which is atropine-sensitive and more commonly observed during immobility, arousal, and anticipation (Bland, 1986; Kramis et al., 1975; Vanderwolf, 1969). Thus, the stationary delay period and active center-arm traversal in our task may engage partially distinct hippocampal network states. In addition, cholinergic tone strongly modulates hippocampal theta dynamics and has been proposed to regulate encoding, retrieval, and memory-guided behavior (Buzsáki, 2002; Hasselmo, 2005). These differences may contribute to the greater behavioral impact of perturbations during active trial progression compared to passive delay periods.

In this view, the opening of the door may represent a critical transition point at which Type I theta is engaged, and stored cue information is transformed into an action plan and organized into a planned behavioral trajectory. Importantly, perturbations delivered later during center-arm traversal, closer to the T-junction, produced smaller impairments, suggesting that the relevant computation may occur early during movement initiation rather than at the final choice point itself. While vicarious trial and error behaviors at the choice point may reflect a subset of trials involving uncertainty or indecision (Redish, 2016), our results suggest that in the majority of trials, the mouse likely commits to a trajectory substantially earlier, well before overt behavioral trajectories diverge. In our task, the critical feature does not appear to be the opening of the door itself or the associated sensory transition, as perturbations delivered 800 ms after door opening, once animals had already initiated movement but remained early in the run, produced similar impairments. Together, these findings suggest that hippocampal-prefrontal circuits may be particularly important during the early formation of a planned behavioral trajectory rather than during sensory processing or final choice execution. This interpretation is consistent with observations that hippocampal and prefrontal circuits represent future goals and spatial targets preferentially during movement and theta-associated states (Johnson and Redish, 2007; Kay et al., 2020; Tanaka et al., 2014; Tang et al., 2025; Vollan et al., 2025; Zutshi et al., 2025).

Overall, our findings suggest that hippocampal and medial prefrontal contributions to memory-guided behavior are dynamically modulated across task epochs, with the strongest requirement emerging during the early implementation of planned actions rather than during passive maintenance alone.

### Limitations and future directions

Several limitations should be considered when interpreting these findings. First, our optogenetic silencing was spatially restricted to dorsal CA1 and dorsal mPFC, and we cannot exclude the possibility that residual activity in untargeted subregions, including ventral hippocampus, which has been specifically implicated in hippocampal-prefrontal communication during encoding (Babl and Sigurdsson, 2025; Spellman et al., 2015), preserved partial function during delay-period manipulations. Second, some perturbation conditions, particularly the longer-delay and delayed-run manipulations, were tested in relatively small cohorts, although effect sizes were consistent with the broader pattern across experiments. Third, our manipulations targeted hippocampus and mPFC independently, and future experiments involving simultaneous perturbations or multi-region recordings will be important for determining how these circuits cooperate across task epochs. Most directly, combining temporally precise silencing with large-scale recordings during navigation would clarify whether hippocampal perturbation during the run epoch disrupts prospective spatial sequences toward a decision, and how that varies temporally within a trial.

## METHODS

### Experimental Animals

This experiment was approved by The Institutional Animal Care and Use Committee at New York University Langone Medical Center. All mice were housed in facilities within the NYU School of Medicine Science Building. Prior to behavioral training, mice were kept in groups of 4-5 per cage under standard environmental conditions: approximately 22°C temperature, ∼45% relative humidity, and a 12-hour light/dark cycle. After behavioral training, the mice underwent surgery and were then placed in single housing and on a reversed 12-hour light cycle, with lights on at 7PM and off at 7AM. While mice had unrestricted access to food throughout, water access was limited during behavioral training to ensure body weight remained at around 80% of their pre-training levels. We used a total of 8 mice (6 male and 2 female) of which three were C57BL/6J wildtype (Jax Stock No. 000664) and five were double transgenic mice crossed between Pvalb-IRES-Cre females (Jax Stock No. 017320) and Ai32 males (Jax Stock No. 024109), and two were also C57BL/6J wildtype but given a Dlx-ChR2-mCherry virus in addition. Mice ranged from ages of 3-6 months old and were 20-30g in weight at the time of implantation.

### Surgical Procedures

Mice were implanted with 200 µm optic fibers to target the CA1 region and/or the prelimbic areas of the medial prefrontal cortex. For all surgical procedures, mice were first induced with 3% isoflurane in oxygen anesthesia (SomnoSuite Low-Flow Anesthesia System) and placed in a stereotaxic apparatus (Kopf Instruments). For the remainder of the surgery, the isoflurane level was lowered to 1.3-1.5%. Throughout the surgery, the mice were placed on a heating pad (Physitemp, TCAT-2LV Animal temperature controller) to maintain body temperature at 37°C. After confirming no reaction to a toe-pinch, Vaseline was applied to the eyes for lubrication, followed by a small incision along the anterior-posterior axis on the scalp. Iodine (Dynarex Povidine Iodine solution) and lidocaine cream (Ferndale LMX4) were applied to the open scalp area, and the skull was scored using a scalpel to ensure proper adhesion to the headcap. The coordinates of the hippocampus (-1.8 mm AP, ±1.5 mm ML from bregma) and medial prefrontal cortex (+2 mm AP, ±0.35 mm ML from bregma) were then marked with a Sharpie marker on the skull. After the base plate of the headcap was cemented to the skull surface using Metabond (Parkell C&B, #S380), a drill (NSK Ultimate XL) was used to make the craniotomy at the coordinates marked by the Sharpie. 200 µm optic fibers were slowly lowered into these holes using clamps to the correct DV coordinates (-0.9 mm for the hippocampus and -1 mm for the mPFC). A combination of mineral oil (Fisher chemical #O121-1) and paraffin wax (Sigma-Aldrich, #18634) (melted in a 1:1 ratio) was used to seal the craniotomy. Light-cured and regular dental cement (Unifast LC, Unifast Trad) was applied around each of the optic fibers to ensure the fibers stay in place. After confirming the optic fibers were securely in place, the clamp was removed. Once all optic fibers were in place, the top portion of the headcap (custom 3D printed, opaque black to prevent light leakage) was cemented to the base plate using dental cement.

Following surgery, an NSAID analgesic was injected (Ketoprofen at 5 mg/kg, ∼0.13 mL, stock solution of 1 mg/mL, subcutaneous). Mice were allowed to recover and were continuously monitored for a week before water deprivation and behavior training resumed.

### Virus Injections

Two out of eight mice were injected with AAV5-mDlx-ChR2-mCherry (titer, 1.4×10^13^, custom prepared from Addgene, plasmids were a gift from Dr. Gord Fishell) in the hippocampus (-1.8 mm AP, ±1.5 mm ML, -1.1 mm DV from bregma, 500 nl volume) and mPFC (+2 mm AP, ±0.35 mm ML, -1.2 mm DV from bregma, 300 nl volume). After each injection, the pipette was left in place for at least 10-15 minutes. The injections were administered at a flow rate between 25-50 nl/min using a microsyringe pump (World Precision Instruments, UMP3 UltraMicroPump). The constructs were allowed to express for at least 2-3 weeks before the mice were implanted with optic fibers.

### Histology

Following the end of experiments and data collection, mice underwent perfusion with 0.9% saline, followed by 4% paraformaldehyde (PFA) in phosphate-buffered saline (PBS) (Affymetrix USB). Extracted brains were immersed in 4% PFA for an additional 24 hours for post-fixation. Coronal brain sections (40-50 µm thick) were then prepared using a vibrating blade microtome (Leica VT1000S) which were then mounted onto electrostatically charged slides, coverslipped with DAPI Fluoromount-G (Southern Biotech, 0100-20). The sections were imaged using a virtual slide microscope (Olympus, VS120). Viral expression and the position of optic fibers were confirmed at the end of each experiment.

### Behavioral apparatus and task description

We constructed a T-maze that consisted of a central stem leading to two perpendicular arms, forming the top of the ‘T’. The track was built using 1/8” thick white acrylic (for the floor and the outer walls) and 1/8” thick transparent acrylic (for the inner walls). The central stem had dimensions of 100 cm (length) × 7 cm (width), with 5.1 cm high walls and each arm had dimensions of 10.8 cm (length) × 7 cm (width), with 5.1 cm high walls. The bottom of this central stem was the ‘home-base’ where the mice started the behavior task behind an initially closed clear door attached to a motor (SunFounder 20KG High Torque Servo Motor SF3218MG). LED strips were placed along the right and left inner walls of the central stem until it reached the junction point. Between each arm of the ‘T’ (right and left) and the central stem, a door attached to a motor was placed. The track contained 3 water ports in total: one at the base of the ‘home-base’ (at the very bottom of the ‘T’) and one at the end of each arm. Water was delivered at these ports through blunt 18G needles connected (using Tygon E-3603 Tubing; 1/16” ID, 1/8” OD) to solenoid valves (Cole-Parmer, EW-98302-02) that were opened for 50 ms. The needle was positioned within a U-shaped infrared (IR) sensor (HiLetgo LM393 Correlation Photoelectric Sensor) to detect the animal’s licks. The same U-shaped IR sensor was attached to the clear door at the ‘home-base’ and was used by the mice to initiate each trial of the behavior task.

Variables such as water delivery, cue/delay duration, door opening, and the trial phase were controlled and coordinated by a custom-made Arduino circuit. For a subset of mice, an overhead camera (FLIR Backfly S) captured images with a frame rate of 60 Hz.

### Delayed cued-choice task

All mice started the behavior task behind the clear door at the ‘home-base’. When the animal nose-poked the IR sensor attached to the clear door and broke the IR beam, a 1-second visual cue was triggered, where either the LED strip on the right or left wall turned on. This was immediately followed by a 1-second delay period (or 3 seconds for the ‘Longer Delay’ task). At the end of the delay period, the clear door opened, and the mice ran the length of the track to turn toward the side of the arm where the cue was presented. It took around 3 seconds for the mouse to reach the junction where it had to decide which arm to enter. If the mice went to the correct side and licked at the water port, the solenoid was triggered, delivering water. The incorrect lick did not trigger the solenoid. After the mice licked at the water port at one of the arms, it returned to the ‘home base’. One final lick at the ‘home-base’ water port reset the trial. The setup of the task, therefore, involved a forced delay (behind the door, 1-s long) and a delay that arose as the mice ran down the track (∼3-s long).

### Behavioral training and analysis

#### Training phase

Prior to implanting the optic fibers, mice were pre-trained to perform the task until they reached around 70% accuracy. The stages of the training included – (1) Habituation to the track and learning to poke water ports for a water reward (∼2-3 days). (2) Forming the stimulus-response association by keeping the door leading to the incorrect arm closed, preventing the mice from entering that arm. The cue stayed on until the mouse poked at the correct water port (∼4-6 days). (3) Confirming if the stimulus-response association was made by having both doors leading to each arm open. The cue stayed on until the mouse poked at the water port at the end of one of the arms. The mice did not move on to the next training stage until they reached around 70-80% accuracy (∼2-6 days). (4) Gradually shortening the cue duration. The mice initially performed the task with long cue durations. If they performed well (∼70-80% accuracy), in the next session, they would perform the task but with a shorter cue duration. The cue duration was gradually shortened until they could perform the task with a 1-second cue (4-10 days). The length of the entire training, from start to finish, varied depending on the mouse. Some mice finished in 8 days while others took 2-3 weeks. The main source of variability was that the mice often developed biases toward a certain arm, which had to be corrected through consecutive trials toward the unpreferred side. Following surgery, mice were retrained from stage (2). To get the mice accustomed to the optic fiber, post-surgery mice were trained on stage (4). Post-surgery mice needed 1-2 weeks to reach 70-90% accuracy on the behavior task.

#### Optogenetic silencing phase

Only one session was performed per day for each mouse, targeting either bilateral hippocampus or mPFC. On each day, optogenetic stimulation was targeted to 1 of 8 phases of the trial. (1) Cue (2) Delay (3) Cue+Delay (4) Run (first 2s) (5) Run (after 1.6s) (6) Run (after 800ms) (7) Delay+Run (8) Run (first 2s) for unilateral silencing. On one of the days, the mice were tested with a longer 3s delay, during which stimulation was targeted to the last 2 seconds of the delay period.

#### Optogenetic stimulation parameters and hardware

Light was delivered by coupling the optic fiber to a laser diode (450 nm blue, Osram) and stimulation pulses were delivered using an isolated current driver (Thorlabs, LDC205C). Light intensity was calibrated across animals and was between 2 mW to 15 mW. Light output was measured using a power meter (Thorlabs, PM100D). Optogenetic stimulation was delivered as a constant square pulse using PulsePal2 (Sanworks, v2), triggered by an Arduino sending TTLs based on the optogenetic targeting section for that session.

#### Video-tracking and position estimation

To estimate the position of the mouse during the task, we used DeepLabCut (version 2.3.9). A subset of frames was manually labeled to mark key body parts (e.g., nose, body center), and a neural network was trained using these labels. The trained network was then used to extract the position of the mouse across all video frames. Only frames with high likelihood estimates (>0.9995) were retained. Tracking data were processed using custom MATLAB scripts and aligned to behavioral events to extract movement trajectories and assess running patterns during different trial phases.

#### Turn Point Detection

Perseveration during stimulation trials prevented the accurate estimation of the point of significant divergence between left and right trials. To estimate the location at which animals committed to a side-arm choice without requiring equal sampling of both sides, we identified the point of maximal lateral deviation along the central arm. For each trial, position traces were smoothed, and the velocity components along the horizontal (x) and forward (y) axes were calculated with respect to time. We then computed the ratio of lateral to forward velocity (dx/dy) as a measure of relative horizontal motion during forward running. The turn point was defined as the y-position at which |dx/dy| was maximal, corresponding to the point of strongest horizontal deviation relative to forward motion.

### Statistical analysis

All statistical tests were performed using MATLAB R2023b. The Wilcoxon signed-rank test was used. Boxplots display the median and 25^th^ and 75^th^ percentiles, and the whiskers show the data range.

### Use of AI-assisted tools

We used ChatGPT (OpenAI) solely for language refinement and minor text editing during manuscript preparation. All scientific content, data interpretation, and conclusions were generated by the authors. The authors reviewed and verified the accuracy of all text and take full responsibility for the final manuscript.

## ACKNOWLEDGEMENTS

We thank R. Kasa for help with behavioral training and Dr. A. Mar for help with the behavioral paradigm design. We also thank the Buzsaki lab for insightful comments throughout the project. Funding: This work has been supported by NIH grants (R01MH122391; U19NS107616) and an NSF grant 1707316 (NeuroNex MINT) to G.B and a Simons Collaboration on the Global Brain Transition to Independence Fellowship to I.Z.

## AUTHOR CONTRIBUTIONS

I.Z., G.B., and T.D. planned and designed the experiments. Experiments were performed by T.D., L.A., and I.Z. Data analysis was performed by T.D., L.A. and I.Z. I.Z and T.D. wrote the paper with input from all authors.

## CONFLICT OF INTERESTS

The authors declare no competing interests.

## Notes

### Competing Interest Statement

The authors have declared no competing interest.

